# *Cylicodiscus gabunensis* (FABACEAE) aqueous stem bark extract mediated silver nanoparticle enhance anti-inflammation on Wistar rats

**DOI:** 10.1101/2024.12.15.628536

**Authors:** Francois Eya’ane Meva, Pamela Nguadie Mponge, Juliette Koube, Gildas Fonye Nyuyfoni, Jean Baptiste Hzounda Fokou, Agnes Antoinette Ntoumba, Thi Hai Yen Beglau, Alex Kevin Tako Djimefo, Annie Guilaine Djuidje, Geordamie Chimi Tchatchouang, Ariane Laure Wounang, Madeleine Ines Danielle Evouna, Maeva Jenna Chameni Nkouankam, Yolaine Pamela Dada Youte, Hassana Moussa Ndotti, Philippe Belle Ebanda Kedi, Sone Enone Bertin, Christoph Janiak

## Abstract

**Introduction:** Plants are a source of bioactive ingredient that can play a key role in development of new drugs. In recent years, plant mediated-biological synthesis of nanoparticles has gained importance due to its simplicity, cost effectiveness and eco-friendly nature. To the best of our readings, nanoparticule from *Cylicodiscus gabunensis* stem bark is still untaped. This work therefore aimed at assessing the anti-inflammatory properties of the silver nanoparticles obtained from the aqueous extract of the stem bark of *Cylicodiscus gabunensis*(Cg).

**Methodology:** *Cylicodiscus gabunensis* extract was prepared by infusion-followed by the biosynthesis of silver nanoparticles. The synthesis was monitored by color and UV-Vis spectrophotometry. Infrared spectroscopy aimed at revealing the functional groups present at the surface of the nanoparticles. Structural elucidation was done by powder X-ray crystallography, while microstructure and elemental mapping was performed with scanning electron microscopy and energy dispersive X-ray spectroscopy. In vitro anti-inflammatory test was done using the BSA denaturation based test. The acute toxicity was done using the OEDC 425 guideline. carrageenan-induced rat paw oedema model was used to ascertain the *in vivo* anti-inflammatory effects.

**Results:** Phytochemical screening of the aqueous extract of *Cylicodiscus gabunensis* revealed the presence of polyphenols, flavonoids, alkaloids, coumarins, saponins, triterpenes, steroids, reducing sugar, tannins, and the absence of anthraquinone. The surface Plasmon resonance peak in the UV-Vis spectrum shows absorption spectra between 380 and 550 nm. Stability studies done over time showed that the nanoparticles were stable even after two months of synthesis. IR spectroscopy revealed the presence of O-H, N-H, C≡C, C=C, C-O, and C=O groups. PXRD confirms the formation of silvernanoparticles (AgNPs) while nanograins of various forms were visualized by SEM and EDX. No toxic sign was observed. The maximum inhibitory percentages were 95% at 200 µg/mL and 91% at 400 μg/Kg for *in vitro* and *in vivo* anti-inflammatory effects, respectively.

**Conclusion:** This paper spot the light on silver nanoparticle from *Cylicodiscus gabunensis* aqueous stem bark for their anti-inflammatory effect on paw oedema model.

## Introduction

*Cylicodiscus gabunensis*, a species of the Fabaceae family, is widespread throughout various regions in Cameroon, including the Babeng locality in the Centre Region, the Lomié forest in the East Region, and at the edges of forests in Mujuka, Loum, and Manjo in the Littoral Region. Ethnobotanical surveys in Cameroon indicate that the stem bark of this plant is used in traditional medicine for the treatment of various ailments such as diarrhea, malaria, bacterial infections, rheumatism, migraine, and stomach pain [1, 2]. Previous studies on the ethyl acetate extract of this plant have revealed the presence of secondary metabolites, as well as its analgesic and anti-inflammatory activities [3].

Inflammation is the response of vascularized tissues to harmful stimuli, such as infectious agents, mechanical damage, and chemical irritants [4]. This inflammatory response can also be triggered by tissue damage or infections, which induce oxidative stress, consequently leading to inflammation [5, 6]. Anti-inflammatory therapy typically employs synthetic molecules, including steroidal (corticosteroids) and nonsteroidal anti-inflammatory drugs. While these drugs are widely used, their side effects can be significant, including immunosuppression and toxicity affecting the renal and digestive systems, respectively [7, 8]. To minimize side effects, natural substances, particularly plants with anti-inflammatory properties, are used in traditional treatments [9]. However, many of these plant extracts hold biologically active constituents that are highly water-soluble and exhibit low absorption due to their high molecular weights. The effectiveness of medicinal plants hinges on the adequate supply of the active substance, highlighting the need for new carriers to deliver these substances at sufficient concentrations directly to the target sites [10].

Silver nanoparticles (AgNPs) have been prepared by new synthesis methods that use biological matter, such as plants or microorganisms, as bioreactors, thereby avoiding the use of non-aqueous solvents [11, 12]. This innovative technology, which organizes metabolites at the metal interface through self-assembly, offers a promising avenue for new drug research. This is particularly relevant for inflammatory pathologies, a field gaining increased attention due to the rising number of deaths caused by inflammatory diseases, which the World Health Organization (WHO) regards as one of the biggest threats to global health [13]. Furthermore, obtaining silver nanoparticles from plants is a guarantee of the preservation of biodiversity due to their significant enhancement of pharmacological actions at low doses [14]. In this light, this research presents the evaluation of the anti-inflammatory effect of silver nanoparticles in the stem bark of *Cylicodiscus gabunensis*.

## Experimental

### Collection, authentication, and preparation of extract

*Cylicodiscus gabunensis* (Cg) stem bark (Figure 1) was harvestedin the locality of Babeng about 4 km from Makak central town, on the road leading to Ngouatè in the Central Region of Cameroon, and authenticated at the Cameroon National Herbarium compared to a voucher specimen previously deposited number 43972/HNC. The fresh stem bark of *C. gabunensis* harvested was cut into small slices, dried in a shade for 3 weeks, and then crushed to powder using a crusher. The aqueous extract of the stem bark powder of *C. gabunensis* (Cg-AE) was obtained by introducing 10 g of the powder into 100 mL of distilled water and heating to 80 °C while maintaining the temperature for 5 minutes using a magnetic stirrer with a heating system. The infusion was then filtered using Whatman filter paper N°1 to get rid of the solid particles [15, 16]. Parts of the aqueous extract obtained were used for the synthesis of silver nanoparticles, for phytochemical screening, and a part was concentrated in the oven at 55 °C for the determination of the extract yield. The dried extract was weighed to assess the extractable content determined by the following formula 1.

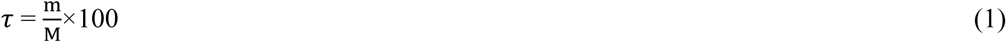

*τ*: extractable content in percent (%);

m: mass of dry extract (g);

M: mass of dry leaves (g);

**Figure 1:**
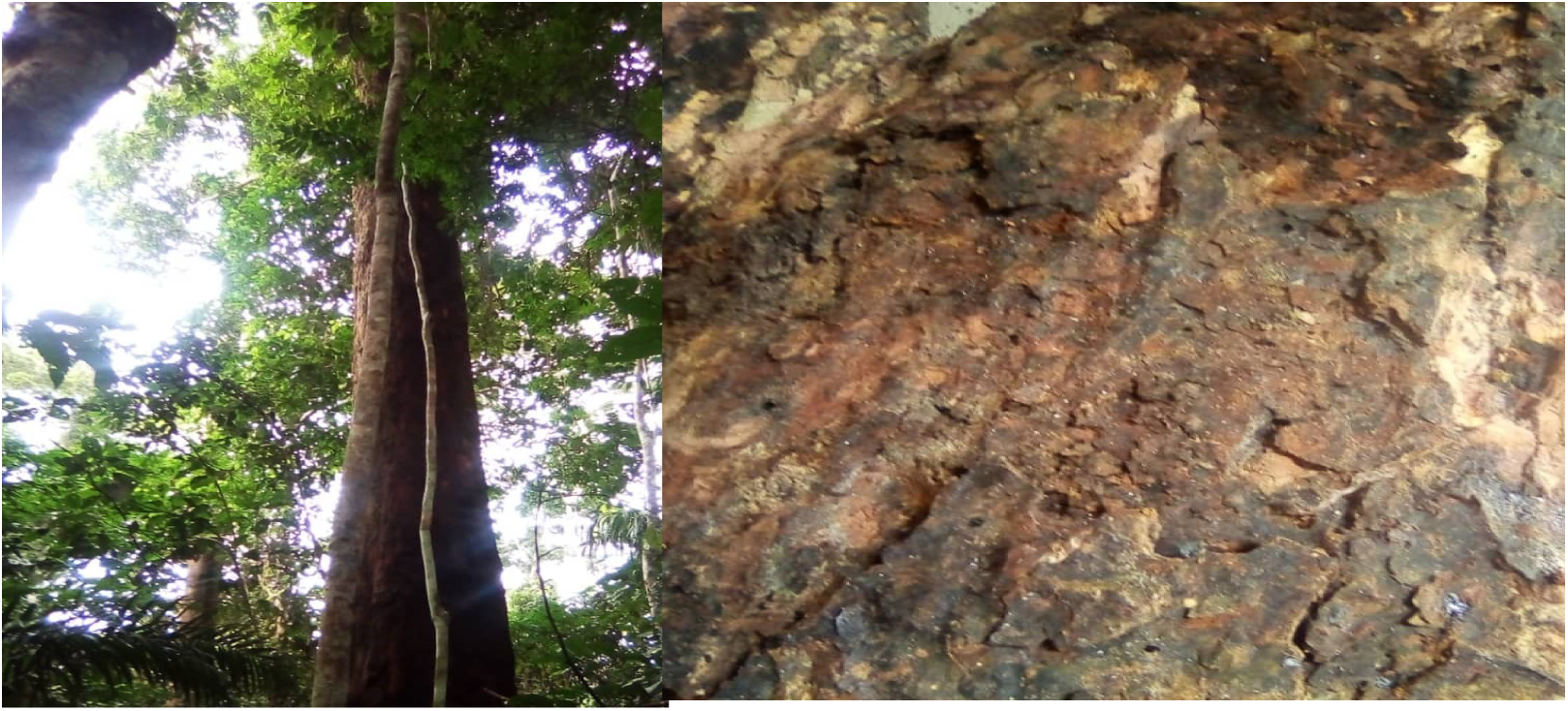
C*y*licodiscus *gabunensis* stem bark.

### Phytochemical screening

The phytochemical screening to detect secondary metabolites alkaloids, flavonoids, terpenoids, coumarins, saponins, anthocyanins, and anthraquinones was performed using the Harborne established protocol [1].

### Synthesis of silver nanoparticles

Silver nanoparticles were obtained as in previous literature [15, 17]. Briefly, 10 mL and 20 mL volumes of Cg-AE were distributed in 6 reaction flasks, and their pH adjusted to 2, 4, 6, 8, 10, and 12 using 0.1 mol/L sulfuric acid and sodium hydroxide, respectively. To these solutions, 50mL of different concentration (10^-1^ ; 10^-2^ and 10^-3^mol/L in water)was added respectively to each tube. This The final solutions were then stored at room temperature and away from light (to avoid the photo- oxidation of silver) [14].

The observation of a color change was the first parameter to assess the formation of silver nanoparticles. Furthermore, 1 mL of the aliquot from each bottle was taken for UV-Vis spectrophotometric scanning, and the absorbance spectra between 350 nm and 900 nm at 5 minutes, 1 hour, 6 hours, 12 hours, 24 hours, 1 week, 2 weeks, and 1 month of incubation were obtained. This was for the search for a surface plasmon resonance characteristic of the formation of AgNPs [14]. The nanoparticles were isolated by centrifugation at 4000 rpm for 1 hour, washed twice with water and ethanol, and finally dried at 50 °Cfor 24hours.

### Ultraviolet-visible characterization (UV-Vis)

Ultraviolet-visible spectroscopy was monitored using an aliquot of 2 mL of suspension on a P9 double beam spectrophotometer from VWR. Measurements were made between 200 and 800 nm.

### Dynamic Light Scattering (DLS)

DLS was performed on a Malvern NanoSZetasizer with a HeNe laser wavelength at 633 nmin water. Three measurements were made per sample.

### Fourier-transformed infrared spectroscopy (FTIR)

Infrared spectroscopy was conducted using a Bruker Tensor 37 with attenuated total reflection (ATR), by scanning between 600 and 4000 cm^-1^.

### Powder X-ray diffraction (PXRD)

The PXRD diffraction of the nanoparticles was performed using a Bruker D2 Phaser powder diffractometer (Cu K-Alpha1 [Å] 1.54060, K-Alpha2 [Å] 1.54443, K-Beta [Å] 1.39225) by preparing a thin film on a low-background silicon sample holder.

### Scanning electron microscopy (SEM) and energy-dispersive X-ray spectroscopy (EDX) investigations

Scanning electron microscopy (SEM) was carried out with a Jeol JSM−6510LV QSEM Advanced electron microscope with a LaB_6_ cathode at 20 kV equipped with a Bruker Xflash 410 (Bruker AXS, Karlsruhe, Germany) silicon drift detector for energy-dispersive X-ray spectrometric (EDX) analysis. The nanoparticle samples were sputtered with gold by using a JEOL JFC-1200 Fine Coater.

#### Animal and ethical considerations

Female albino rats (Rattusnorvegicus) aged 8-12 weeks and weighing 120-180 g, were grown in the Animal Facility Laboratory of the Faculty of Medicine and Pharmaceutical Sciences of the University of Douala, Cameroon. The animals were sorted randomly in standard polypropylene cages in groups of three and maintained under standard conditions of temperature (24 ± 2 °C) and light (approximately 12h light/dark cycle) with free access to standard laboratory diet and tap water *ad libitum*.

### Evaluation of acute toxicity of the aqueous extract and silver nanoparticles of *C. gabunensis*

The acute toxicity test was carried out according to guideline 425 of the Organization for Economic Cooperation and Development (OECD), which is the limit test at the dose of 2000 mg/kg body weight (b.w) of rats of the Wistar strain [18]. Three groups of 3 female rats were used; two groups received 2000 mg/Kg b.w of *C. gabunensis* aqueous extract and silver nanoparticles respectively and the other batch which served as the control group received 10 mL/Kg b.w of distilled water. Rats were repartitioned evenly based on their masses before being subjected without food but not water 12 hours before the test and weighed before the administration of the samples.

Observation of clinical signs was done after 30 minutes, 1, 2, and 4 hours after administration of the samples and distilled water. 4 hours later the rats were hydrated and fed. The observation of the clinical signs was carried out every day for 14 days. The following clinical signs were observed: modification in the skin; hair on the rats; nature of their eyes; presence or absence of trembling; convulsions; salivation; diarrhea; sleep.

Similarly, the mass of the rats was taken every day for the first 2 days and later, taken after 2 days and the growth rate calculated. After 14 days, the rats were sacrificed by ether aspiration. The organs (heart, lung, liver, kidney, and spleen) removed after dissection, were rinsed with 0.9% saline solution, then weighed.

### Evaluation of the anti-inflammatory activity of the aqueous extract and silver nanoparticles of *Cylicodiscusgabunensis*

#### The Bovine Serum Albumin (BSA) denaturation test

The Albumin denaturation test was used to put in place the anti-inflammatory property of the silver nanoparticles from *C. gabunensis*aqueous stem bark following the protocol described in Belle *et al.,* with slight modification [17]. A 5 mL rational mixture constituting 0.2 mL of 1 % BSA solution, 2.8 mL PBS (pH= 6.4) and 2 mL of variable concentrations of the aqueous extract and the silver nanoparticles (25, 50, 100, 150 and 200 μg/mL). The same volume of distilled water served as the negative control, and diclofenac at the same concentrations of 25, 50, 100, 150 and 200 μg/mLserved as the positive control (reference drug). The 5 mL rational mixture was then incubated at 37 °C for 15- 20 minutes and then heated to 70 °C for 5 minutes. After cooling, the absorbance was measured at 600 nm using the UV-Vis spectrophotometer. The percentage inhibition of the denaturation of albumin was calculated using the formula 2:

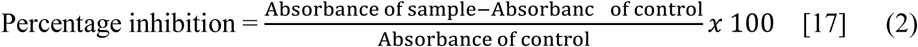

#### The Carrageenan-induced rat paw edema method

The standard protocol was described by Winter *et al* in 1962 [18] with slight modifications. Rats were randomly divided into 6 groups of 5 rats each and treated as follows: Group 1: 10 mL/Kg b.w of distilled water, (Control);Group 2: 10 mg/Kg b.w of diclofenac, (Standard);Group 3: 100 µg/Kg b.w of Cg-AgNPs, (test 1);Group 4: 200 µg/Kg b.w of Cg-AgNPs, (test 2);Group 5: 400 µg/Kg b.w of Cg- AgNPs, (test 3);Group 6: 200 mg/Kg b.w of Cg-AE, (test 4).Inflammation was induced by sub-plantar injection of 0.1 mL of carrageenan (1% carrageenan suspended in 0.9% NaCl) in the right hind paw of each rat. The injection was made one hour following oral administration of the various substances (distilled water, diclofenac, silver nanoparticles, and the aqueous extract). Measurement of paw size was done before carrageenan injection and 30 min, 1, 2, 3, 4, 5, and 6 hours after the carrageenan injection using a digital calliper. The anti-inflammatory activity was evaluated as percentage inhibition of oedema in each treated group compared to control using the formula3:

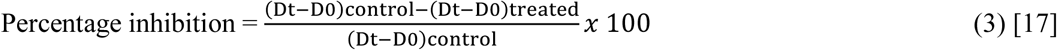

Dt= average diameter for each group after treatment

D0= average diameter for each group before treatment

### Statistical analysis

Excel software was used to plot the UV-Vis spectra. This software also helped us to record all data obtained during experiments. Results were expressed as the mean ± SEM. The difference between treated groups and control groups was compared using two-way analysis of variance (ANOVA), followed by Dunnett’s post hoc test for pairwise comparisons. The statistical analysis was performed using GraphPad Prism Software version 8.01, and the probability values less than 0.05 were considered statistically significant.

## Results

Synthesis and characterization of silver nanoparticles of the aqueous extract of *C. gabunensis*

### Extraction yield

After an aqueous extraction by infusion of 25 g of *C. gabunensis* stem bark powder in 250 mL distilled water and concentrated in an oven at 55 °C, a brown powder weighing 4.09 g was obtained corresponding to an extraction rate of 6 %.

### Phytochemical screening of the aqueous extract of *C. gabunensis*

Table 1 presents the results of the phytochemical screening of the aqueous extract of the stem bark of *C. gabunensis* and the supernatant obtained after centrifuging the solution of silver nanoparticles.The phytochemical screening of Cg-AE revealed the presence of polyphenols, phenols, flavonoids, alkaloids, coumarins, saponins, triterpenes, steroids, reducing sugars and tannins and the absence of anthraquinone.

**Table 1.** Phytochemical screening of plant extract and supernatant.

| Secondary metabolites | Aqueous extract | Supernatant |
| --- | --- | --- |
| Polyphenols and phenols | + | - |
| Flavonoids | + | - |
| Alcaloides | + | - |
| Coumarins | + | - |
| Saponins | + | + |
| Triterpenes | + | - |
| Steroides | + | - |
| Reducingsugar | + | - |
| Anthraquinone | - | - |
| Tanins | + | - |

(+) present; (-) absent

The phytochemical screening of the supernatant of Cg-AgNPs revealed the presence of saponins and the absence of polyphenols, phenols, flavonoids, alkaloids, coumarins, triterpenes, steroids, reducing sugar and tannins and the absence of anthraquinone.

### Synthesis of silver nanoparticles with the aqueous extract of *C. gabunensis*

The first parameter to confirm the formation of nanoparticles is the visual observation of a color change. Figure 2 presents the change in color after the addition of silver nitrate solution to the aqueous extract of *C. gabunensis*. The mixture of the aqueous extract (B) of *C. gabunensis* (orange color) with a 10^-1^ mol/Lsilver nitrate solution (colorless) (A) gave a brown color (C) after 5 minutes of incubation thereby indicating the formation of silver nanoparticles.

**Figure 2:**
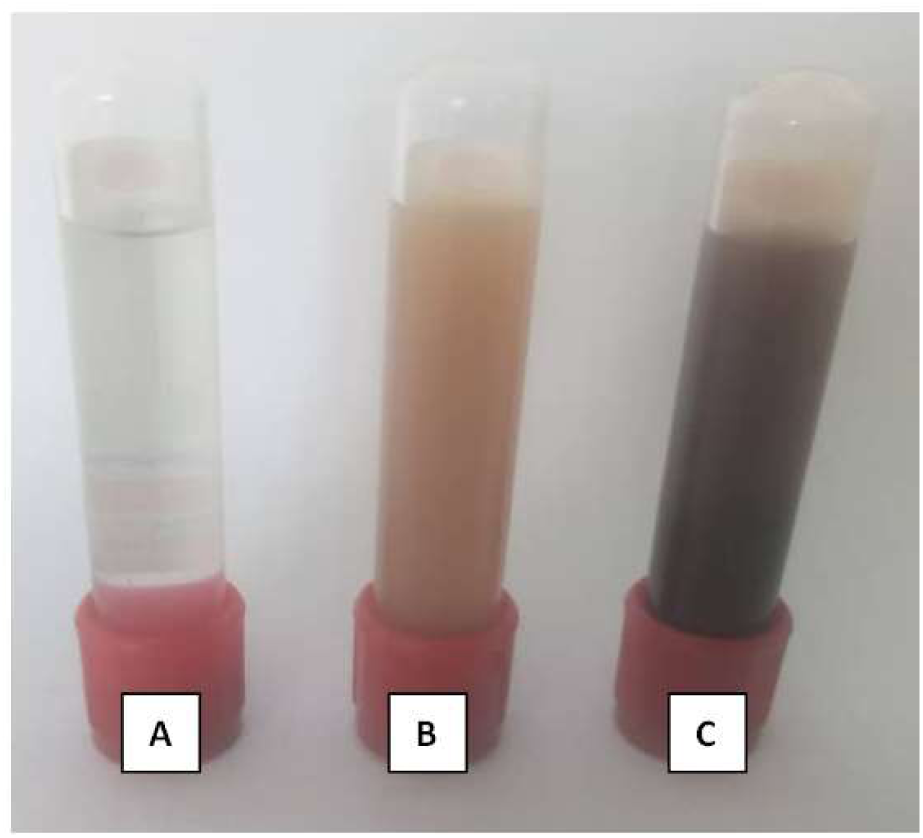
Silver nitrate solution (A), *C. gabunensis* aqueous extract (B) and *C. gabunensis* silver nanoparticles (C)

#### UV-Vis spectral analysis

UV-Vis was used to monitor the bio-reduction of silver ions by the constituents present in the aqueous extract. Figure 3a presents the contact time study. The absorbance spectra were measured between 5 minutes and 2 weeks for a mixture of 2 mL aqueous *C. gabunensis* extract to 10 mL of 10^-1^ mol/L silver nitrate solution. The spectra show the appearance of a broad band of the surface plasmon resonance of AgNPs as from the 5^th^ minute between 380 and 550 nm. The intensity of the absorption band increases with time and reaches a maximum at 4 days with an absorption intensity of 2.1. A shift of the maximum to higher wavelength can be observed after the 1^st^ hour related to formation of bigger particles.

**Figure 3a:**
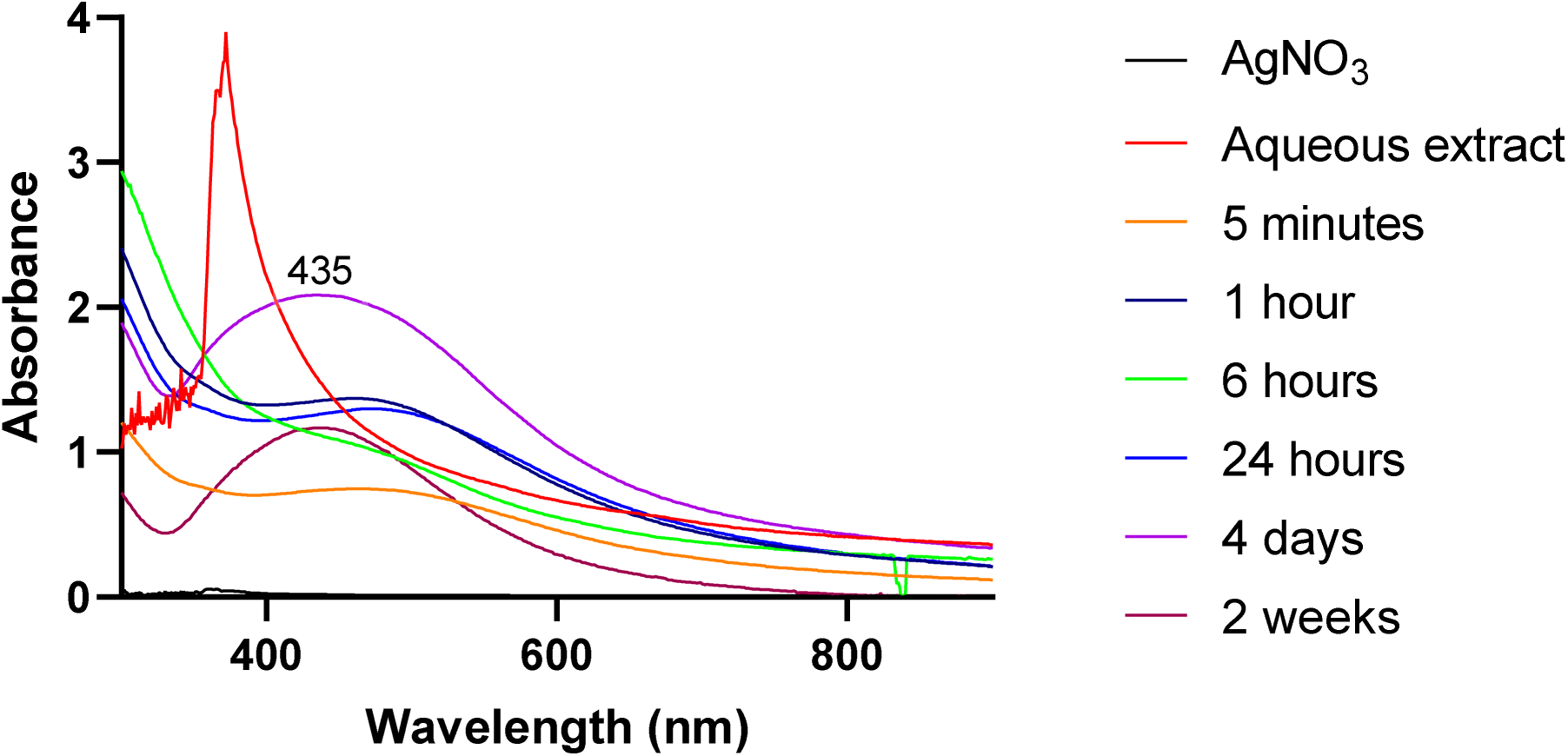
UV-Vis spectrum of AgNPs as a function of contact time.

Figures 3b shows an increase in the intensity of the plasmonic resonance band proportional to the increase in pH of the extract. Secondary metabolites including polyphenols at basic pH have a higher rate of reduction of Ag^+^ and stabilization of Ag^0^ [19]. A clear Plasmon resonance is obtained at pH 10 comparatively to shoulders at other pH.

**Figure 3b:**
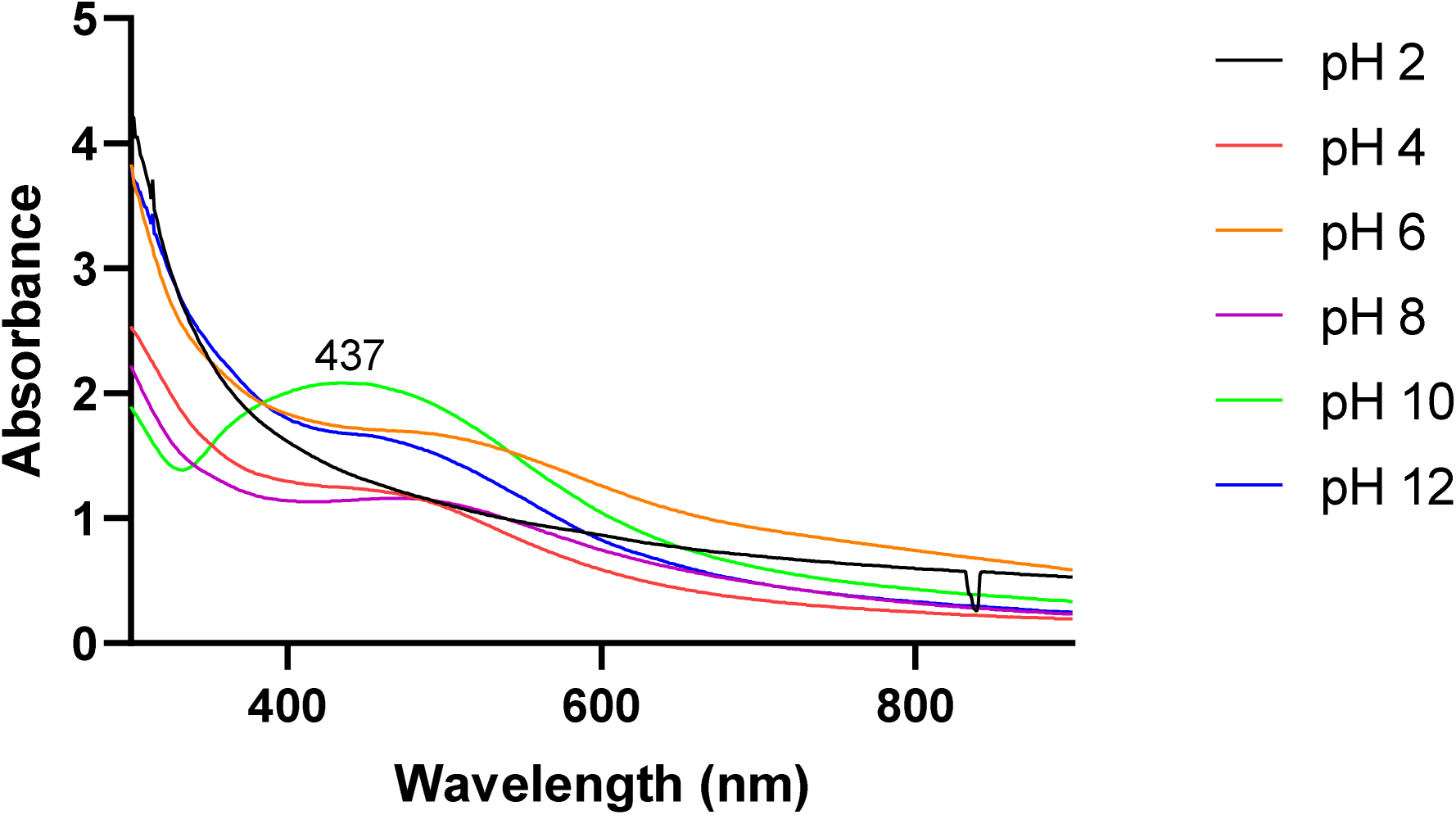
UV-Vis of AgNPs as a function of pH.

Figures 3c present an increase in the intensity of the plasmonic resonance band proportional to the extract’s volume due to increase density of nanoparticles in solution.

**Figure 3c:**
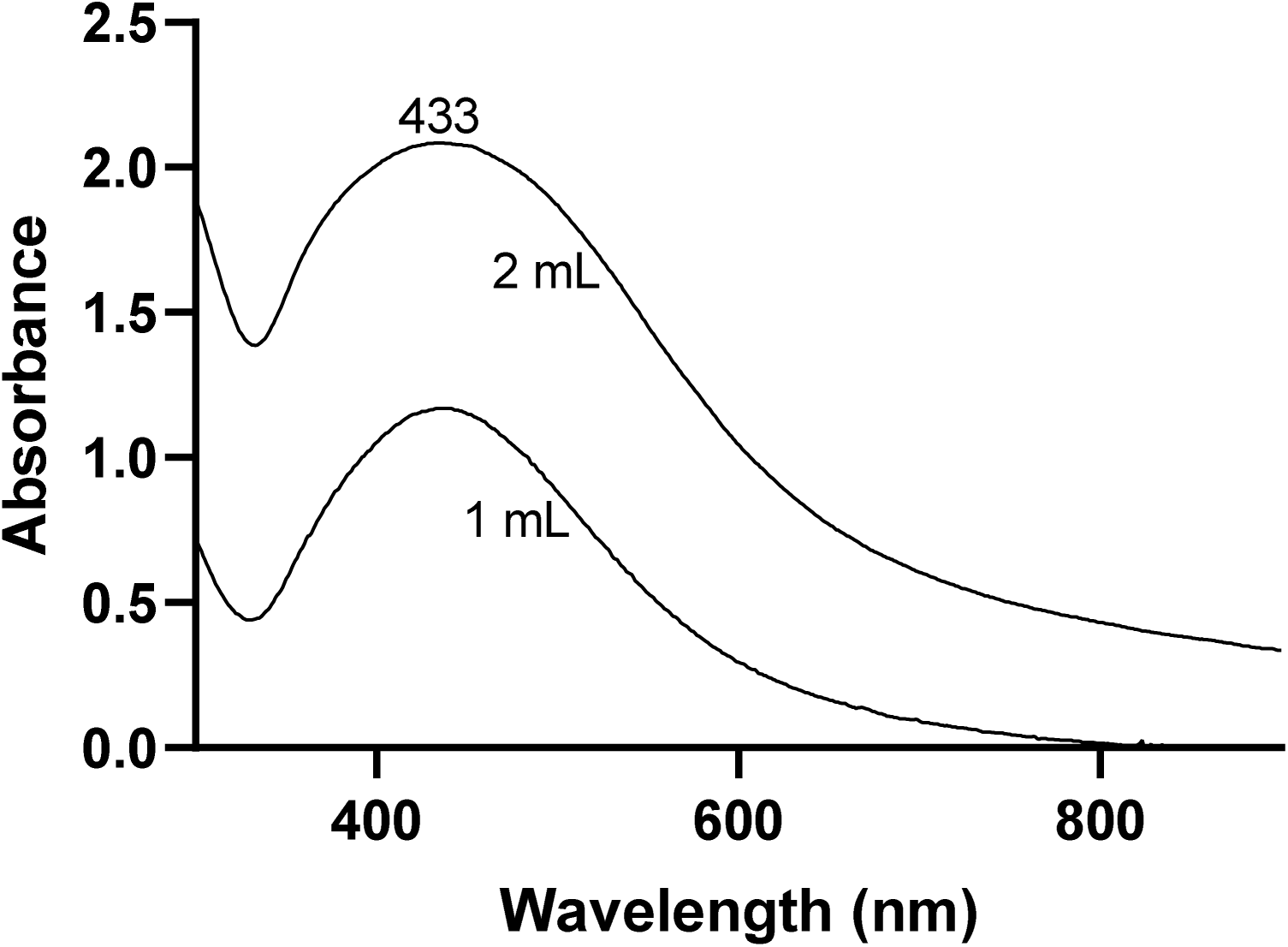
UV-Vis spectrum of AgNPs as a function of extract volume.

Figures 3d presents an increase in the intensity of the plasmonic resonance band proportional to the increase in the concentration of silver nitrate solution due to the availability of free Ag+ to react with secondary metabolites.

**Figure 3d:**
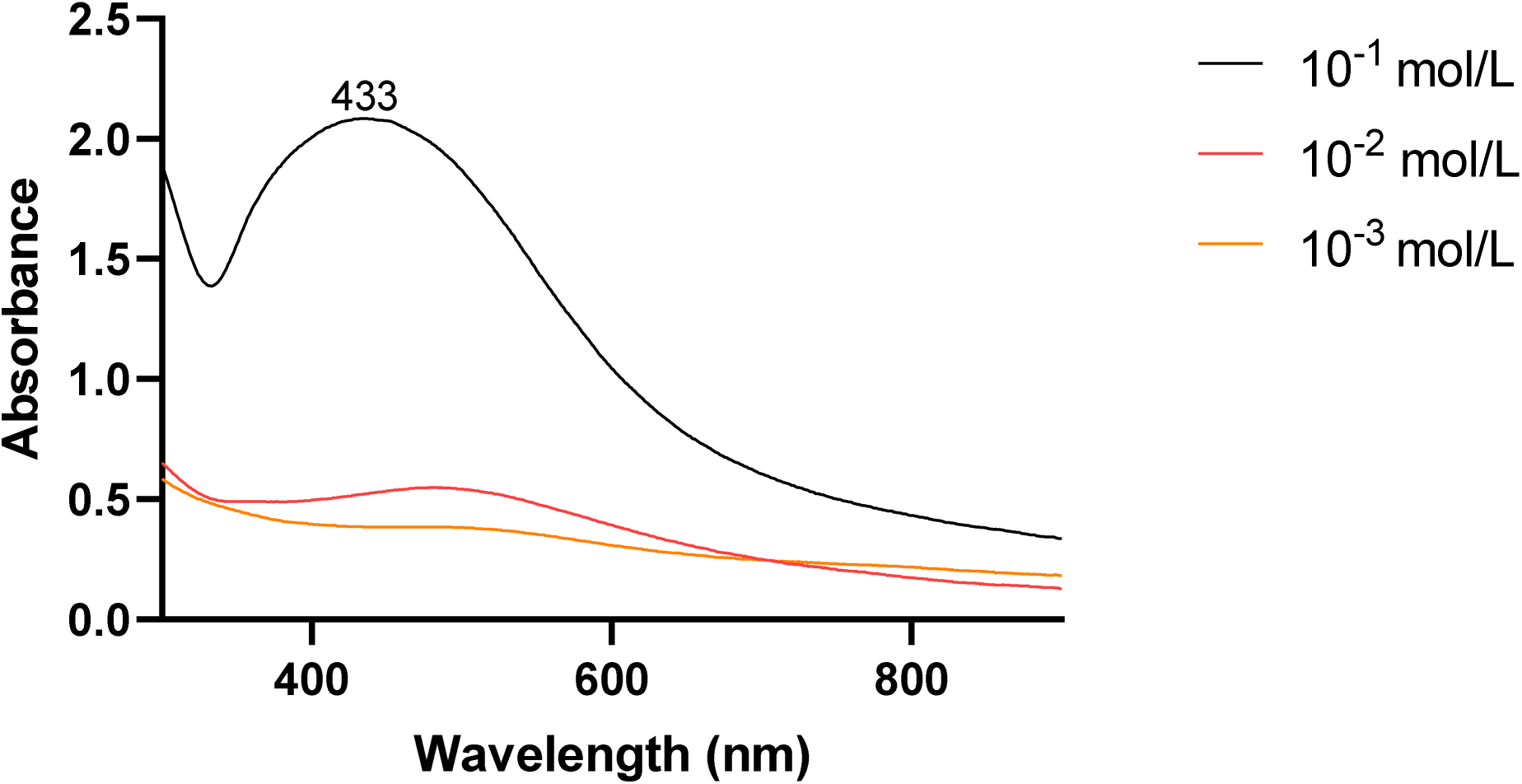
UV-Vis spectrum of AgNPs as a function of silver nitrate concentration.

### FTIR Spectroscopy

IR spectral analysis was realized on the AgNPs powder between 4000 cm^-1^ and 450 cm^-1^. Figure 4 represents the IR spectrum registered which revealed the different functional groups available at the surface of the AgNPs.The interactions between the chemical moieties present in the AgNPs as analyzed by IR spectroscopy, revealed the main vibrations (stretching and bending) in the mentioned wavelength range.

Figure 4 (Table 2) shows in the spectrum of AgNPs the presence of a peak at 669 cm^-1^ and 541 cm^-1^ linked to the vibrations of Ag-O stretching; a high intensity peak at 3264 cm^-1^ linked to the vibrations of an asymmetric stretch N-H bonds, O-H of amine, alcohol compounds, and H-bonded water respectively; an additional peak of intensity at 2914/2844 cm^-1^ characteristic of the vibrations of symmetric stretch of the CH_2_/CH_3_bonds; another peak of intensity at 1610 cm^-1^characteristic of the vibrations of asymmetric stretch of the carbonyl bond confirmed by the presence of the band at 1020 cm^-1^ characteristic of C-O bond.

**Figure 4:**
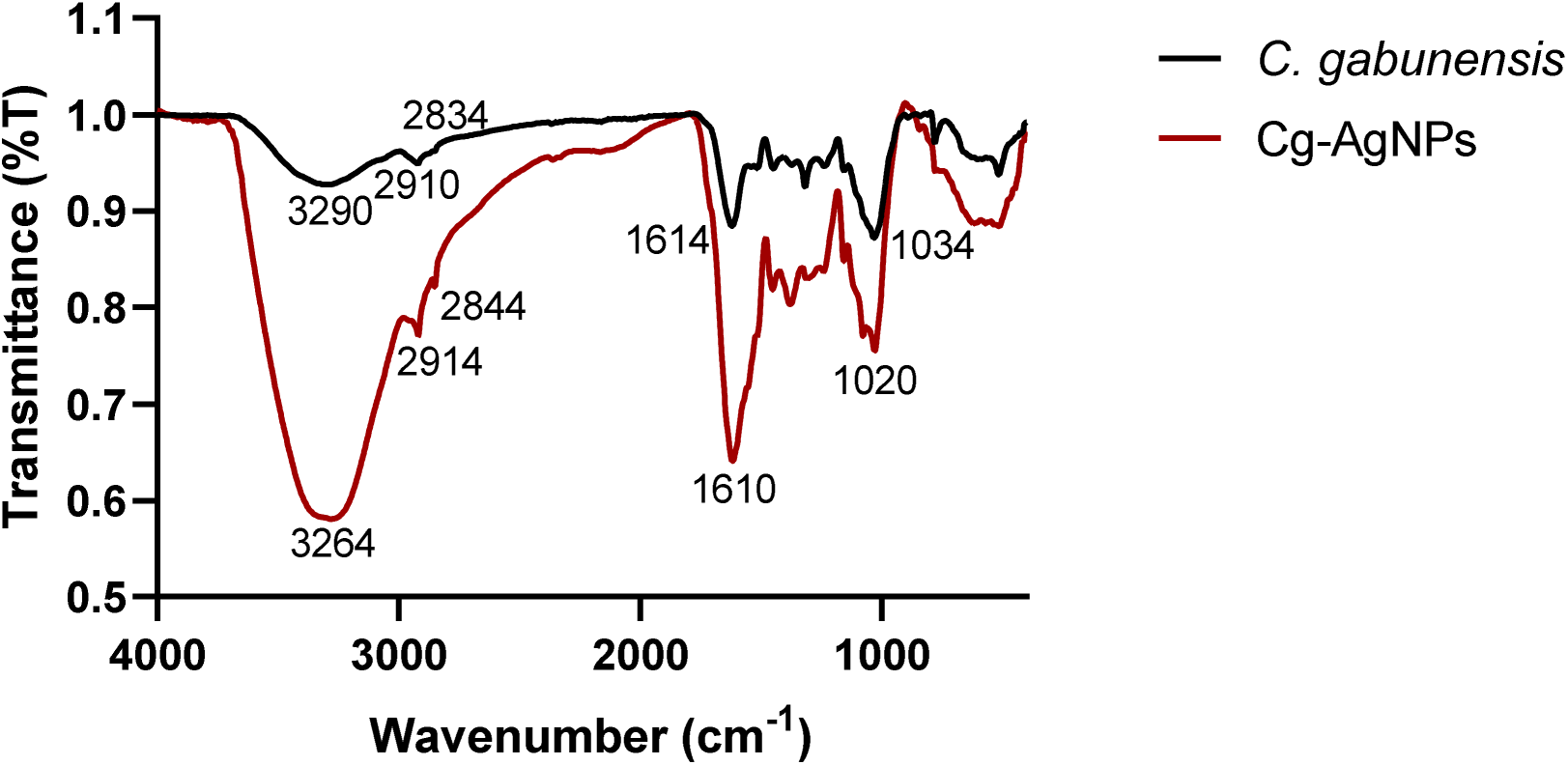
IR spectra of dried *C. gabunensis* extract and Cg-AgNPs.

**Table 2:**
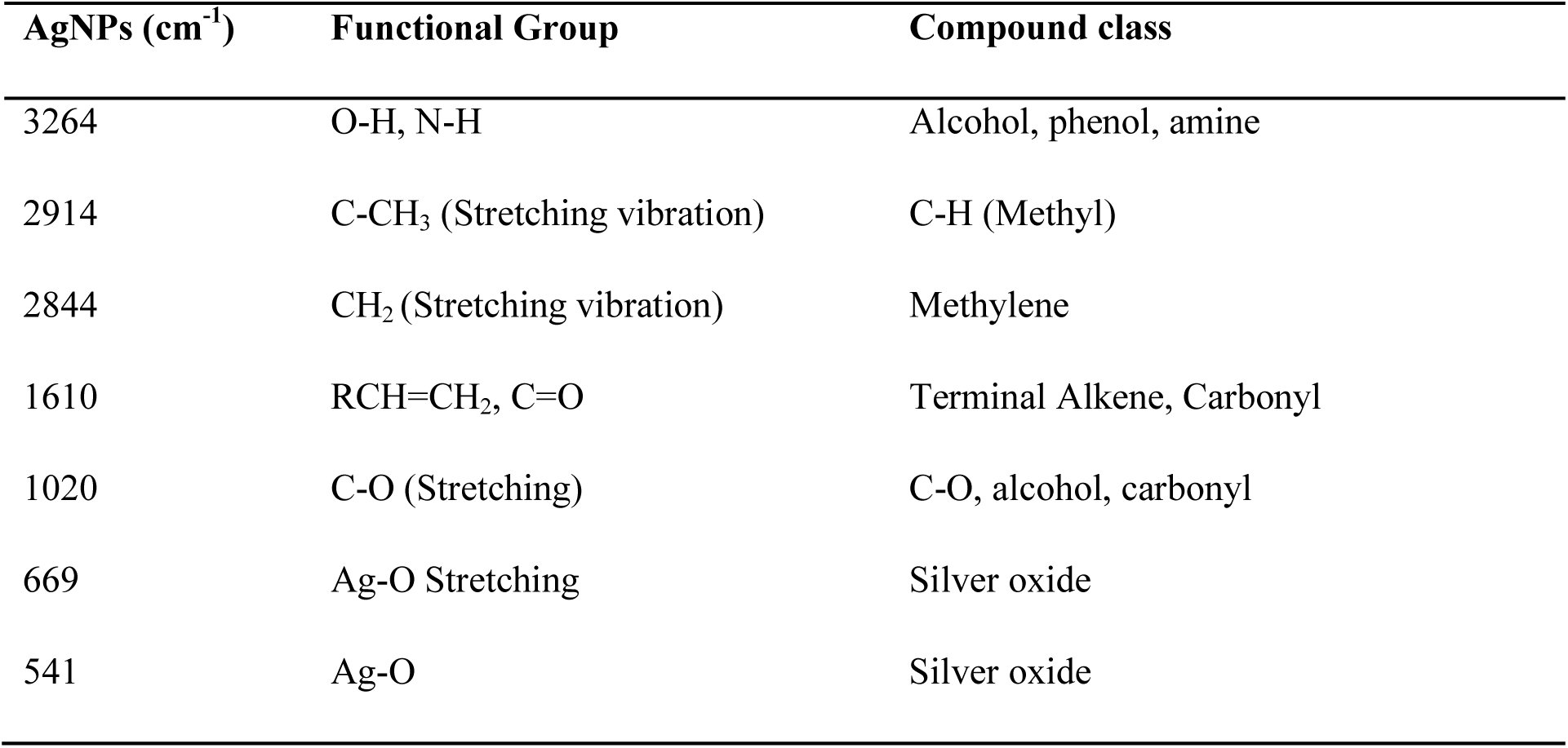
Functional groups obtained from the IR spectra of AgNPs.

| AgNPs ($\text{cm}^{-1}$) | Functional Group | Compound class |
| --- | --- | --- |
| 3264 | O-H, N-H | Alcohol, phenol, amine |
| 2914 | C-CH<sub>3</sub> (Stretching vibration) | C-H (Methyl) |
| 2844 | CH<sub>2</sub> (Stretching vibration) | Methylene |
| 1610 | RCH=CH<sub>2</sub>, C=O | Terminal Alkene, Carbonyl |
| 1020 | C-O (Stretching) | C-O, alcohol, carbonyl |
| 669 | Ag-O Stretching | Silver oxide |
| 541 | Ag-O | Silver oxide |

### PXRD

PXRD was used to confirm the crystalline nature of the AgNPs. The PXRD pattern of AgNPs synthesized from fresh *C. gabunensis* extractis shown in Figure 5. The indexing process of the powder diffraction pattern is performed, and Miller indices (hkl) to each peak are assigned. Four peaks with 2θ values (between 20 and 80 °C) of 38°, 44°, 64°, and 77°2 theta correspond to the (111), (200), (220), and (311) planes of silver. These peaks were observed and compared to the International Centre for Diffraction Data ICDD 04-0783. The data was recorded between 20 and 80°. Size of 10 nm were obtained by using the Scherrer equation with the most intense signal (111).

**Figure 5:**
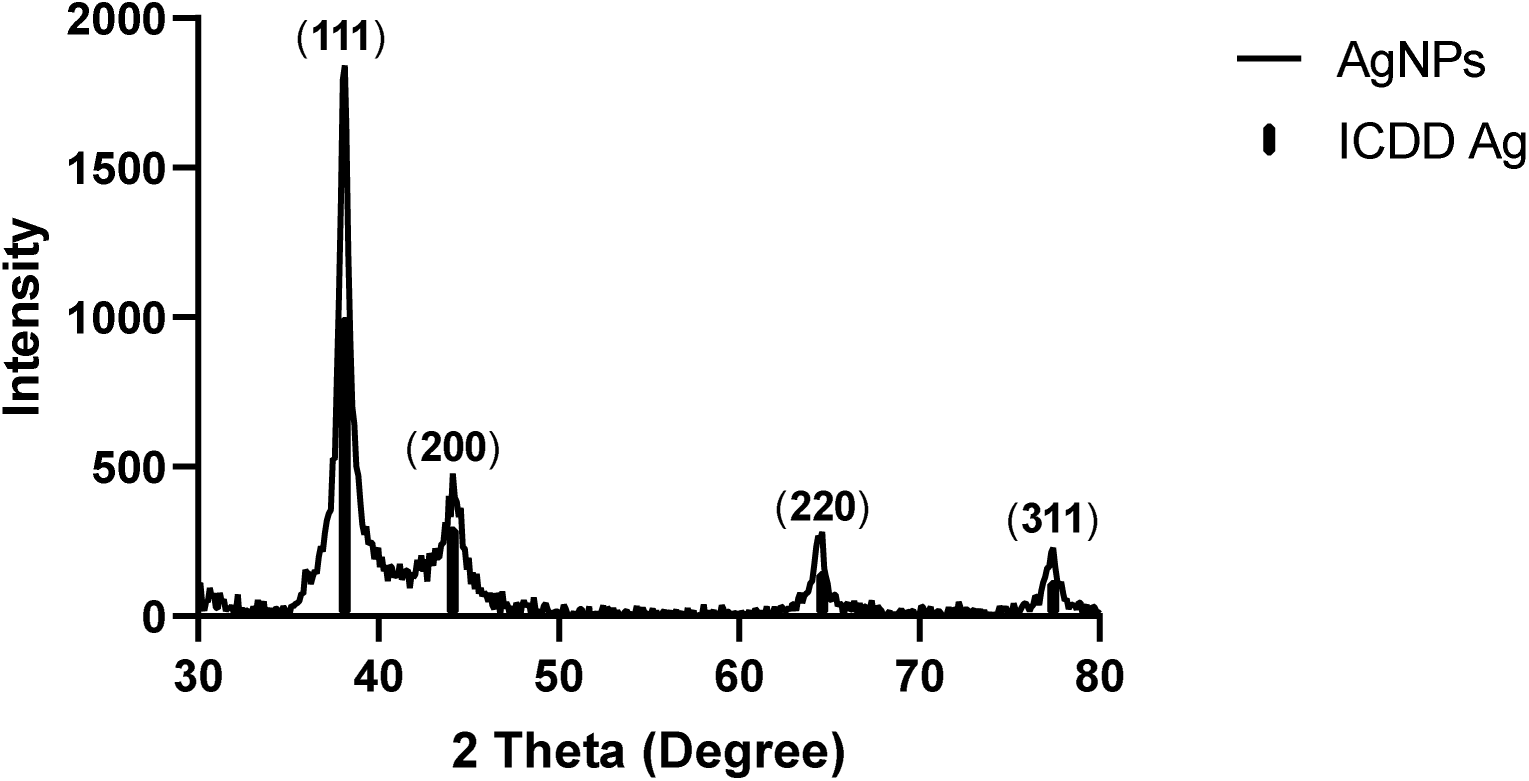
PXRDof AgNPs from *C.gabunensis* and bar diffractogram of AgICDD 04-0783.

### SEM and EDX

SEM assessed the morphology and shows spherical agglomerated particles from the reaction of AgNO_3_ with the *C. gabunensis* extract after separation by centrifugation and drying (Figure 6). Elemental characterization by EDX shows a major composition of O, C and Ag (Figure 7).

**Figure 6.**
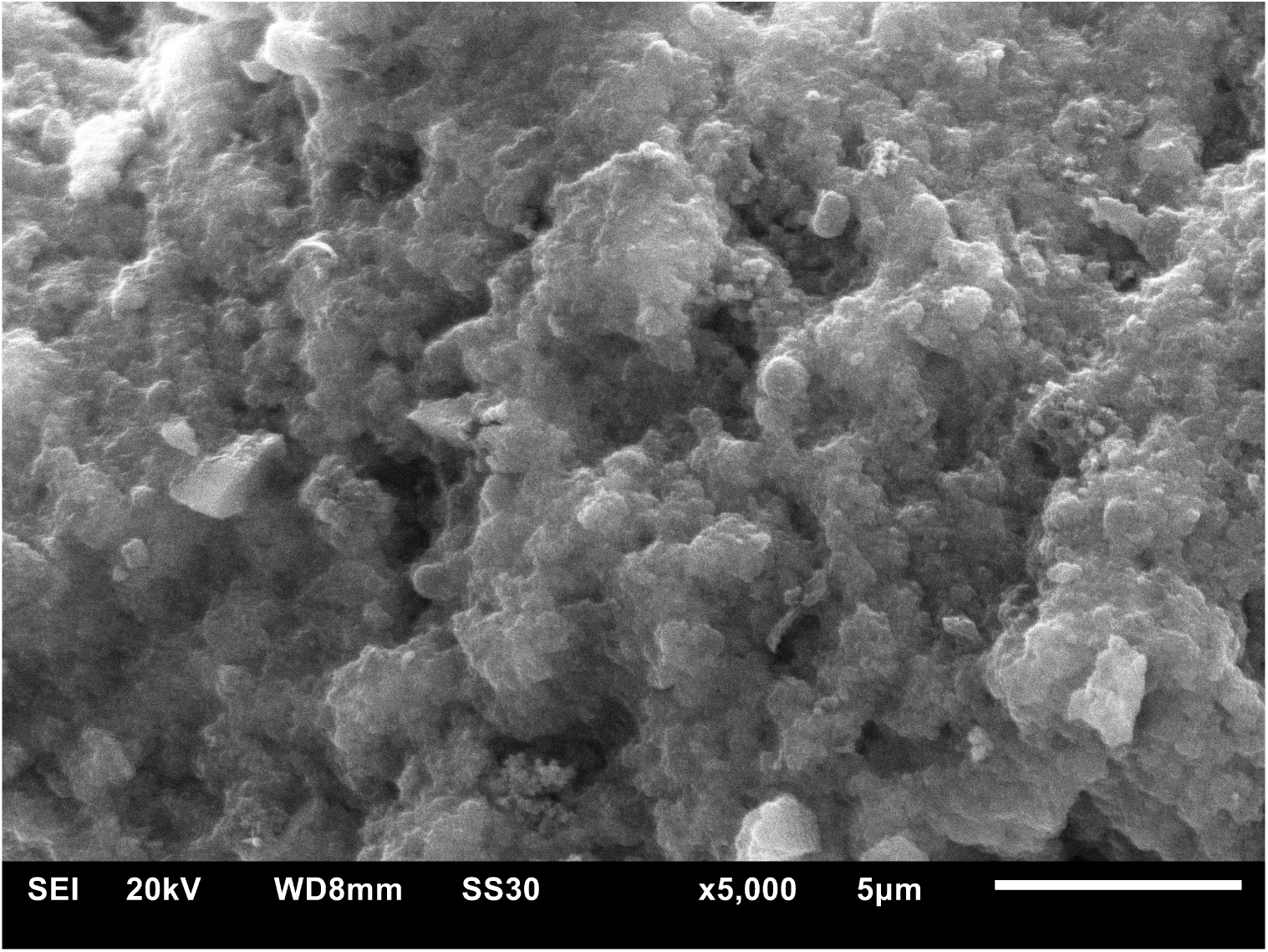
SEM image of AgNPs.

**Figure 7.**
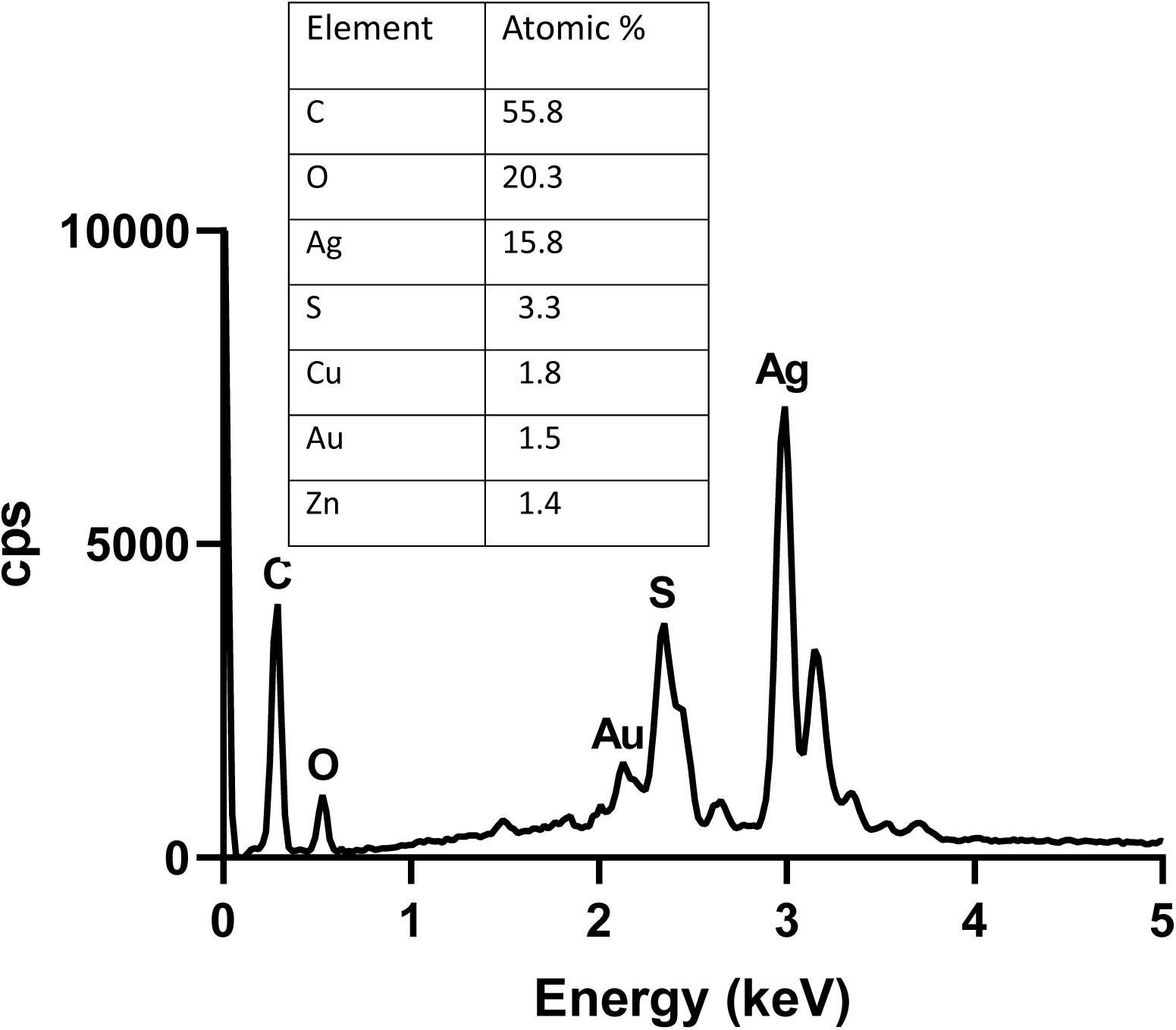
X-ray energy dispersive spectroscopy and element mapping.

Acute toxicological profile of silver nanoparticles of the aqueous extract of *C. gabunensis*

Effect of silver nanoparticles of the aqueous extract of *C. gabunensis* on the clinical parameters in rats

Table 3 shows the clinical parameters of the rats after administration of the aqueous extract and the silver nanoparticles of the aqueous extract of *C. gabunensis*at the limit dose of 2000 mg/kg b.w. During this period of 14 days, no deaths were registered, and rats showed no signs of toxicity.

**Table 3:** Effect of the aqueous extract and silver nanoparticles of *C. gabunensis* on the clinical parameters of rats submitted to acute toxicity over a period of 14 days.

| Clinical signs | Water<br>(1 mL/100 g) | Cg-AE<br>(2000 mg/Kg) | Cg-AgNPs<br>(2000 mg/kg) |
| --- | --- | --- | --- |
| Hair modification | A | A | A |
| Eye modification | A | A | A |
| Trembling | A | A | A |
| Convulsion | A | A | A |
| Stool texture | N | N | N |
| **Salivation** | A | A | A |
| **Mobility alteration** | A | A | A |

A: absent; N: normal

Effect of silver nanoparticles of the aqueous extract of *C. gabunensis* on the variation in the weight of the rats during the toxicity period

Figure 8 presents the evolution in the weight of the rats submitted to acute toxicity. It reveals a normal increase in the weights of the rats with no significant difference between the control group and the test groups for probability values greater than 0.05.

**Figure 8.**
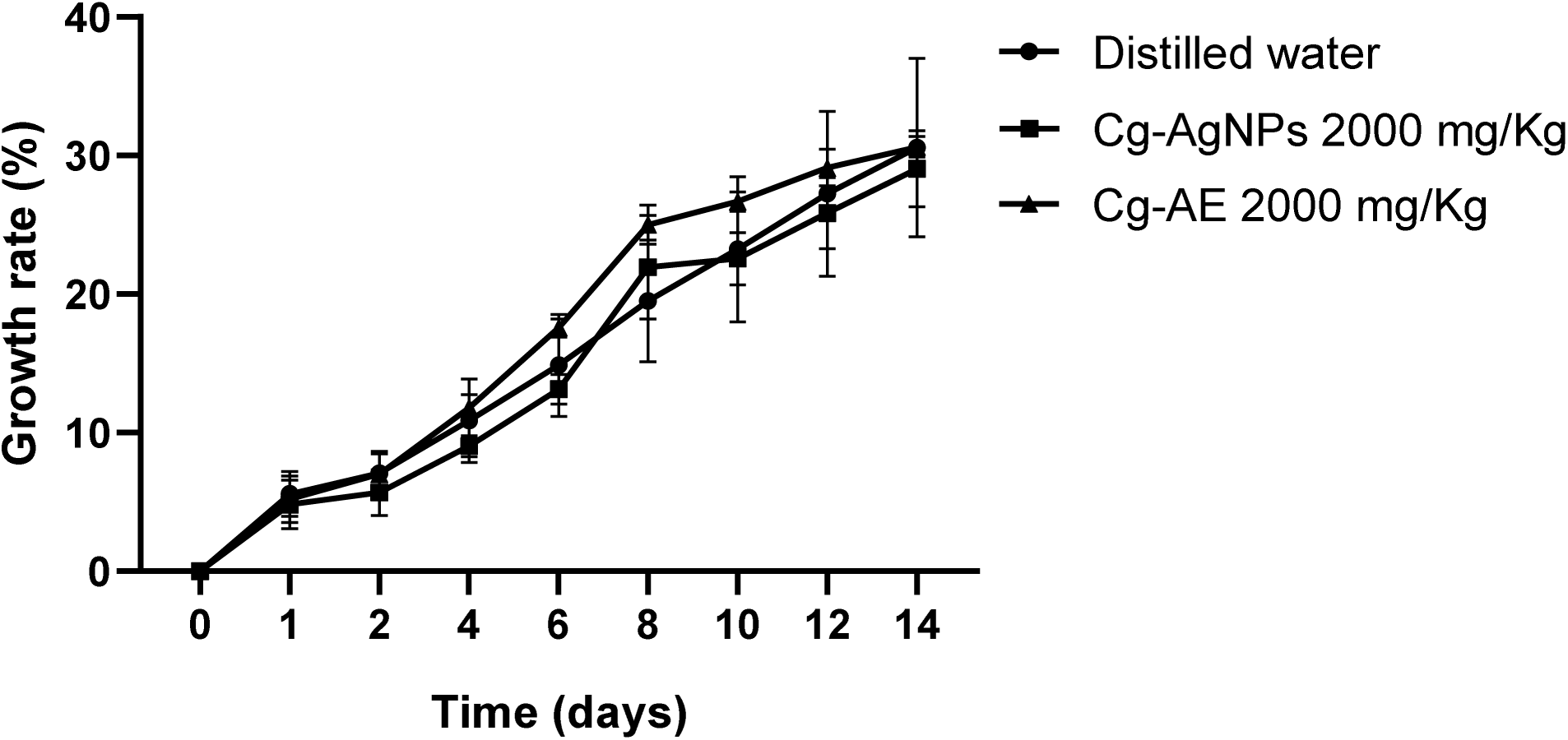
Effect of aqueous extract and silver nanoparticles of *C. gabunensis* on the evolution of the weight of rats over a period of 14 days.

### Effect of silver nanoparticles of the aqueous extract of *C. gabunensis*on the weights of some organs of the rats

Figure 9 shows a comparative analysis of the relative weights of organs of the rats of the test group to those of the control group. It presents the relative weights of organs, and the level of significant differences determined statistically. A slight increase in the weight of the liver of the group treated with Cg-AE 2000 mg/Kg b.w and that of the group treated with Cg-AgNPs2000 mg/Kg b.w was observed with a significant difference for p values less than 0.05 (p= 0.0113) of the test groups to the control group.

**Figure 9.**
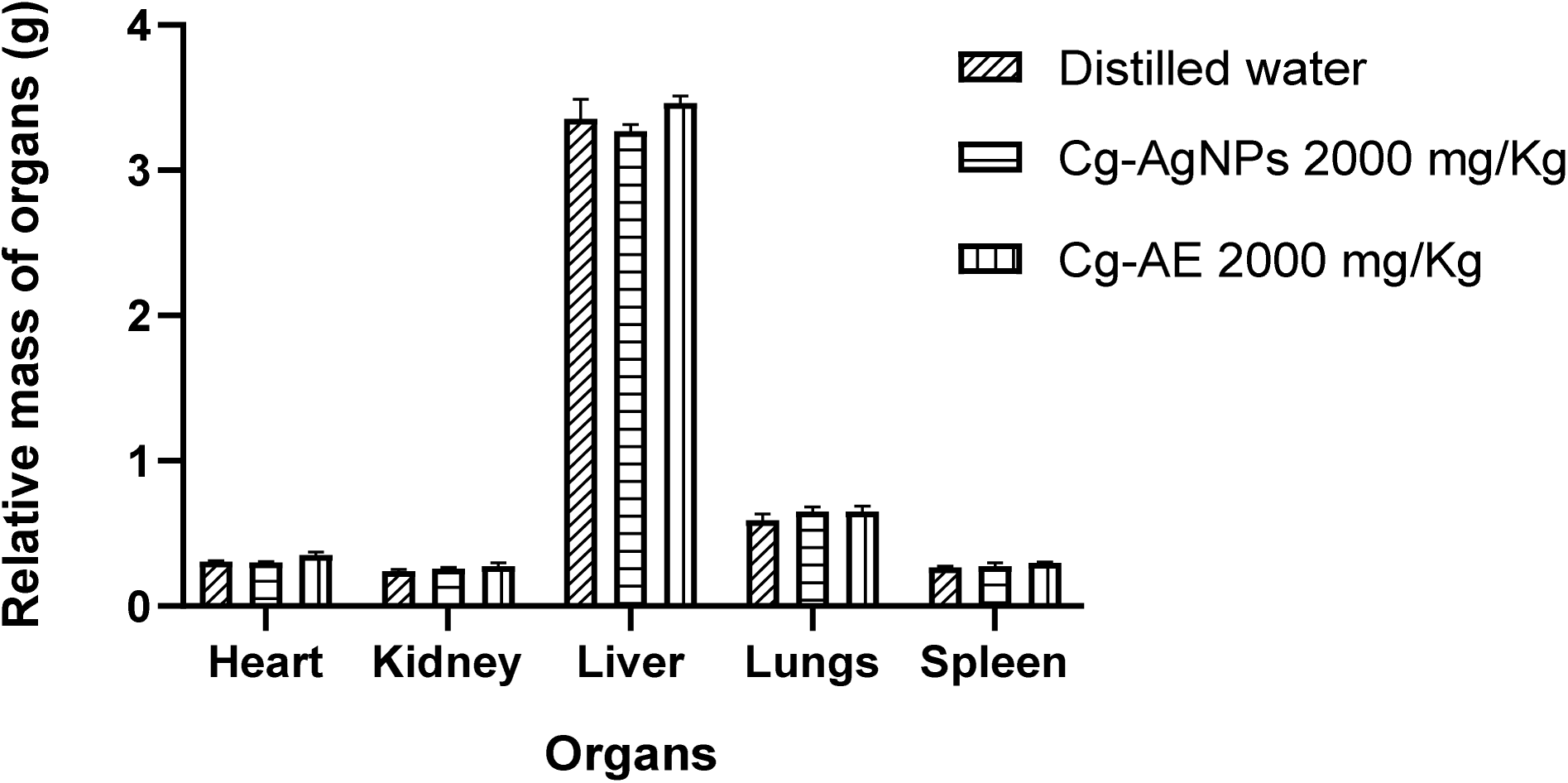
Effect of silver nanoparticles of the aqueous extract of *C. gabunensis*on the weights of organs of rats after toxicity test.

### Anti-inflammatory effect of silver nanoparticles of the aqueous extract of *C. gabunensis In vitro* anti-inflammatory study

Table 4 summarizes the *in vitro* bioassay results of the anti-inflammatory effect of Cg-AE and Cg- AgNPs assessed against heat-induced BSA denaturation. A significant increase in inhibition in a concentration-dependent manner was observed in the test groups and the group that received diclofenac compared to the control group. The maximum inhibition percentage was 76 % for Cg-AE and 95 % for Cg-AgNPs at the highest tested concentration (200 µg/mL) while diclofenac, used as a standard drug, exhibited an inhibition of 58 % at the same concentration.

**Table 4.** Effect of silver nanoparticles of the aqueous extract of *C. gabunensis* on BSA denaturation.

| Sample | Concentration ($\mu\text{g/ml}$) | Optical density ($\pm\text{SEM}$) | % inhibition |
| --- | --- | --- | --- |
| Control | — | 2.98±0.02 | — |
| Cg- AgNPs | 25 | 0.93±0.01<sup>d</sup> | 69 |
| Cg-AgNPs | 50 | 0.73±0.02<sup>d</sup> | 76 |
| Cg- AgNPs | 100 | 0.53±0.01<sup>d</sup> | 82 |
| Cg- AgNPs | 150 | 0.34±0.02<sup>d</sup> | 87 |
| Cg-AgNPs | 200 | 0.15±0.01<sup>d</sup> | 95 |
| **Cg-AE** | 25 | 2.00±0.02<sup>d</sup> | 33 |
| **Cg-AE** | 50 | 1.6±0.4<sup>d</sup> | 47 |
| **Cg-AE** | 100 | 1.09±0.05<sup>d</sup> | 63 |
| **Cg-AE** | 150 | 0.09±0.03<sup>d</sup> | 69 |
| **Cg-AE** | 200 | 0.73±0.02 | 76 |
| **Diclofénac** | 25 | 2.02±0.02<sup>d</sup> | 32 |
| **Diclofénac** | 50 | 1.78±0.04<sup>d</sup> | 40 |
| **Diclofénac** | 100 | 1.36±0.01<sup>d</sup> | 47 |
| **Diclofénac** | 150 | 1.36±0.03<sup>c</sup> | 54 |
| **Diclofenac** | 200 | 1.25±0.02<sup>c</sup> | 58 |

Values are expressed as means ± SEM in each test group (n=3); values having no letter are significantly identical compared with the control, a = p <0.05, b = p <0.01, c = p <0.001, d = p <0.0001.

### *In vivo* anti-inflammatory study

Table 5 shows the evolution of edema and the percentage inhibition in the groups with respect to time. A comparative analysis carried out shows that the injection of carrageenan produced local edema within 30 min after administration, Maximum edema was observed 30 minutes later for Cg-AgNPs and Cg-AE and 1 hour later for the diclofenac. A reduction in edema was appreciated progressively up to the 6^th^ hour, with maximum inhibitory effects of 85 % respectively by Cg-AgNPs at 0.4 mg/Kg b.w of the rat. Diclofenac showed maximum inhibition of 66 % 5 hours after carrageenan administration. The results obtained are all significantly different amongst and in comparison, with the negative control group.

**Table 5.** Effect of silver nanoparticles of the aqueous extract of *Cylicodiscus gabunensis* on rat hind paw edema.

| **Treatment** | **Dose (mg/kg)** | **Diameter ± SEM (cm) (% inhibition)** |  |  |  |  |  |  |
| --- | --- | --- | --- | --- | --- | --- | --- | --- |
|  |  | 30 min | 1 h | 2 h | 3 h | 4h | 5 h | 6 h |
| **Control** | — | 0.69 ± 0.07 | 0.74 ± 0.03 | 0.77 ± 0.04 | 0.7 ± 0.6 | 0.60 ± 0.04 | 0.54 ± 0.07 | 0.5 ± 0.7 |
| **Diclofenac** | 10 | 0.65±0.02<sup>d</sup><br>(20) | 0.7±0.2<sup>d</sup><br>(23) | 0.62±0.04<sup>d</sup><br>(46) | 0.56±0.0<sup>d</sup><br>(43) | 0.50±0.04<sup>d</sup><br>(58) | 0.46±0.03<sup>d</sup><br>(66) | 0.51±0.02<sup>a</sup><br>(-4) |
| **Cg-AgNPs** | 0.1 | 0.75±0.09<sup>d</sup><br>(-11) | 0.68±0.03<sup>d</sup><br>(26) | 0.66±0.03<sup>d</sup><br>(38) | 0.50±0.05<sup>d</sup><br>(71) | 0.50±0.01<sup>b</sup><br>(64) | 0.51±0.02<sup>b</sup><br>(40) | 0.47±0.08<br>(49) |
| **Cg-AgNPs** | 0.2 | 0.6±0.4<sup>d</sup><br>(21) | 0.60±0.02<sup>d</sup><br>(40) | 0.57±0.05<sup>d</sup><br>(54) | 0.5±0.3<sup>d</sup><br>(56) | 0.43±0.07<sup>d</sup><br>(87) | 0.42±0.02<sup>d</sup><br>(83) | 0.44±0.02<sup>d</sup><br>(64) |
| **Cg-AgNPs** | 0.4 | 0.59±0.05<sup>d</sup><br>(46) | 0.58±0.07<sup>d</sup><br>(56) | 0.52±0.05<sup>d</sup><br>(76) | 0.49±0.02<sup>d</sup><br>(76) | 0.45±0.02<sup>d</sup><br>(90) | 0.44±0.04<sup>d</sup><br>(91) | 0.44±0.03<sup>d</sup><br>(85) |
| **Cg-AE** | 400 | 0.70±0.04<sup>a</sup><br>(-0.3) | 0.68±0.03<sup>d</sup><br>(20) | 0.61±0.01<sup>d</sup><br>(46) | 0.52±0.05<sup>b</sup><br>(58) | 0.57±0.02<sup>d</sup><br>(18) | 0.4±0.1<sup>d</sup><br>(29) | 0.4±0.1<sup>d</sup><br>(76) |

Values are expressed as means ± SEM in each group (n=5); values having no letter are significantly identical compared with the control, a = p <0.05, b = p <0.01, c = p <0.001, d = p <0.0001. Values within parentheses are the percentage of inhibition. The negative values show that the drug action has reduced and therefore there is a decrease in the diameter of the hind paw of the negative control group compared to that of the treated group in question.

## Discussion

The aqueous extraction of the dry stem bark powder of *Cylicodiscus gabunensis* gave a percentage yield of 6 %. On comparing this percentage yield with that obtained by Bayoi [19] which was 4 %, it is suggested that the extraction yield is linked to the aqueous extraction method used (decoction), the time of harvest and the efficiency of the filtration process during extraction. Phytochemical screening of the aqueous extract of *C. gabunensis* revealed the presence of polyphenols, phenols, flavonoids, alkaloids, coumarins, saponins, triterpenes, steroids, reducing sugar and tannins. Anthraquinone was absent. This result is coherent with studies carried out by Eya’ane Meva *et al*. [20]. These phytochemicals present in the stem bark of this plant can have a link to the pharmacological properties theypossess such as anti-malaria, anti-psoriasis, anti-rheumatism, antimigraine, gastro protective, and antidiabetic [19]. The absence of secondary metabolites in synthetic supernatant media suggests that the metabolites have been used to reduce Ag^+^ and to stabilize the Ag^0^ nanoparticles. The marked presence of diverse metabolites such as alkaloids, polyphenols, flavonoids, triterpenes, coumarins and tannins is compatible with their anti-inflammatory effects and can be responsible for the reduction of Ag^+^ toAgNPs and their stabilization [12].

The formation of AgNPs was observed after 5 minutes of incubation via a color change. The change in color is due to the reduction of Ag^+^ to Ag^0^ by the secondary metabolites. More precisely, the change in color would occur due to the vibration of the free electrons at the nanoparticle surface [17]. The formation of the nanoparticles was confirmed using the UV-Vis spectrophotometer, by obtaining a surface plasmon resonance band between 380 and 550 nm. Band’s significant width is linked to the large distribution of nanoparticles’ size [21]. The AgNPs are formed from the 5^th^ minute of incubation and their formation continues up to four days of incubation, when it reaches the peak and remains stable for two weeks. This stability is due to the engulfing nature of the ligands made up of secondary metabolites around nanoparticles [17]. The evolution of the formation of AgNPs was observed under various conditions namely; pH, the silver nitrate concentration, the volume of extract and the reaction time (figure 3). The best conditions of synthesis and good stability of Cg-AgNPs were pH 10, silver nitrate concentration of 10^-1^ M, 1 mL volume of extract and 4 days of reaction. It is known that basic pH speed upthe formations of silver nanoparticles [21]. Functional groups present on the surface of the nanoparticles were revealed byFTIR.These functional groups align with the findings of Zhu *et al*. [22], who suggested that phenolic compounds play a role in the reduction and stabilization of nanoparticles. Nanoparticles of pure silver were obtained based PXRD with few nanometers size, comparable to nanoparticles obtained with leaf extract of *Pedalium murex* [23]. The nanograins present aggregates in the solid state with silver based elemental mapping.

The result obtained after evaluation of the acute toxicity of silver nanoparticles of *C. gabunensis* stem bark aqueous extracts shows the absence of toxicity. This result suggests that the lethal dose (LD_50_) of Cg-AgNPs is greater than the limit dose administered (2000 mg/Kg b.w). This result correlates to those obtained by Tchangou Njiemou *et al.* [15] which showed that nanoparticles are not toxic at the dose of 2000 mg/Kg b.w.

Previous studies have evidence that the denaturation of proteins, in particular blood proteins, is a major cause of rheumatoid arthritis and inflammatory [24]. Protein denaturation is a process in which proteins lose their secondary and tertiary structure by application of external physical or chemical constraints such as heat. It is well known that proteins lose their biological functions upon denaturation [25].

The inflammatory condition induced by the sub-plantar injection of carrageenan in rats involves the stepwise release of vasoactive substances such as serotonin, histamine and kinins in the early phase (0- 2 h) and prostaglandins and cyclooxygenase products in the late acute phase (>3 h) [26]. These chemical substances increase the vascular permeability upon their production, thereby promoting the accumulation of fluid in tissues that account for the oedema [27]. Oral pre-treatment of animals with Cg-AgNPs and Cg-AE resulted in an effective inhibition of edema showing that silver nanoparticles have a higher activity compared to that of the aqueous extract and the standard drug. Moreover, this result suggests that synthesized silver nanoparticles from *C. gabunensis* aqueous stem bark extract silver nanoparticles may reduce/inhibit the release of acute inflammatory mediators [17]. The anti-inflammatory properties exhibited by the Cg-AgNPs is due to the individual or synergistic action of various phytochemicals coated to the nanoparticles.

## Conclusion

This study highlights the synthesis of small silver nanoparticles from *C. gabunensis* aqueous stem bark extracts with anti-inflammatory potential. The nanoparticles wear metabolites at the interface and lead to a safe solid aggregate on rat models. This suggests that these nanoparticles can be recommended for formulation of new antioedemateous drug.

## Ethics Statement

The animals were examined and adapted to the new environmental conditions for a week before the formal experiment. All experimental procedures were in strict compliance with theapproved protocol by the Institutional Ethic Committee For Human Research of the University of Douala (Protocol approvalnumber 3144IEC-UD/06/2022/T).

## Acknowledgements

FEM thank the DAAD for a generous Visiting Professor Fellowship (grant no. 57588364).

## Data availability

The authors declare that the data supporting the findings of this study including raw data files are available from the corresponding author upon reasonable request.

## Statement on animal rights

On behalf of all authors, the corresponding author affirms that animal rights were upheld in the study

## Declarations

All authors approve the submission

## Author Contribution

Francois Eya’ane Meva, Christoph Janiak; Writing – review & editing, Conceptualization, Supervision, Investigation, Funding acquisition. Pamela Nguadie Mponge, Juliette Koube, Gildas Fonye Nyuyfoni, Jean Baptiste Hzounda Fokou, Agnes Antoinette Ntoumba; Writing – review & editing, Conceptualization, Visualization. Thi Hai Yen Beglau, Alex Kevin Tako Djimefo, Annie Guilaine Djuidje, Geordamie Chimi Tchatchouang, Ariane Laure Wounang, Madeleine Ines Danielle Evouna, Maeva Jenna Chameni Nkouankam, Yolaine Pamela Dada Youte, Hassana Moussa Ndotti Writing – review & editing, Investigation, Data curation, Animal models, Instrumental support. Philippe Belle Ebanda Kedi, Sone Enone Bertin; Writing – original draft, Investigation, Data curation, Animal models, Writing – review & editing, Investigation,

